# A Tissue Proteolysis Activity Mapping and Substrate Discovery Platform for Identifying Novel Tumor-Activated Biosensors

**DOI:** 10.1101/2025.08.18.670718

**Authors:** Itay Algov, Audrey Van Heest, Megan Theresa Hopton, Frances Liang, Aidan Holmes, Liangliang Hao, Xin Zhou

**Affiliations:** Department of Cancer Biology, Dana-Farber Cancer Institute, Boston, MA, USA; Department of Biological Chemistry and Molecular Pharmacology, Harvard Medical School, Boston, MA, USA; Department of Biomedical Engineering, Boston University, Boston, MA, USA; Biological Design Center, Boston University, Boston, MA, USA

## Abstract

Dysregulated extracellular proteolytic activity is a prominent hallmark of cancer and can thus be exploited for tumor detection and therapeutic development. However, the discovery of tumor-responsive probes has been hindered by the lack of methods capable of capturing proteolytic events directly in tissue samples. Here, we report PSurf, a platform that enables the identification of tissue-specific protease sensors with tissue specimens. Through differential selection of tumor-specific sequences over healthy tissue, PSurf identified context-specific tumor-activated probes that precisely distinguish metastatic lesions in lung tissue slices. Using these substrates, we engineered nanobody-targeted biosensors that release urinary reporters upon tumor-specific cleavage *in vivo*, enabling precise noninvasive tumor detection in a murine lung metastasis model. PSurf provides a foundation for developing conditionally activated agents through tissue-specific activity mapping and probe discovery.

## Introduction

Dysregulation of extracellular tumor microenvironment (TME) proteases is a well-established hallmark of cancer, driving fundamental processes such as angiogenesis, tumor growth, and metastasis. These hydrolytic enzymes play a pivotal role in shaping the TME by mediating matrix degradation, receptor activation, and immune signaling, thereby influencing tumor progression and therapeutic response within a specific disease ecosystem. Harnessing protease activities to sense and respond to specific TMEs has therefore emerged as a promising strategy for detecting and treating disease with high precision^1^. Local proteolytic activation has been utilized to develop imaging agents^2-5^ and activity-based diagnostics^6^ for early cancer detection and treatment monitoring. In addition, conditional activation of therapeutics^7^ can aid the balance between potency and toxicity^8, 9^. This design principle has been implemented to create prodrugs^10^, pro-antibodies, pro-cytokines^11^ and antibody-drug conjugates^12, 13^, several of which have advanced to regulatory approval or clinical trials^14, 15^. Tailored proteolytic profiling holds promise to guide clinical intervention for tumor metastasis, the leading cause of cancer mortality, through the development of activatable tools to precisely detect, monitor, and treat malignant lesions.

Although significant progress has been made over the past decade in characterizing tumor-associated proteolytic activities, most studies have focused on identifying substrates of individual proteases selected based on transcriptomic data^16-20^. The human proteome includes over 500 proteases^21^, yet active recombinant forms are available for only a small subset, limiting the scope of recombinant enzyme-based profiling^22^. Moreover, individual enzyme-based approaches cannot capture the integrated activities occurring within the TME, where diverse proteases function simultaneously and are tightly regulated *via* various activation or inhibitory mechanisms such as cofactors, prodomains, and natural inhibitors such as tissue inhibitors of metalloproteinases (TIMPs)^23-25^. As a result, recombinant enzyme activity may not accurately reflect total proteolytic activity in a tissue context, where the number of target enzymes and regulatory components is much higher. Additionally, synthetic peptide libraries are constrained by limited size and high costs associated with scaling, with most libraries containing only hundreds of peptides^26-32^. More recently, phage display libraries and mass spectrometry-based techniques have advanced protease research, but these approaches remain confined to individual enzyme characterization or are focused on studying native cleavage sequences rather than discovering new substrates^28, 33-43^. To date, no existing method allows direct profiling of proteolytic activity and substrate discovery with living tissue samples.

Here, we report a tissue proteolytic mapping and substrate discovery platform termed PSurf (<u>P</u>roteolysis <u>S</u>ignature discovery via yeast s<u>urf</u>ace display) (**Fig. 1a**). Distinct from acellular platforms, PSurf harnesses genetically encoded yeast surface display coupled with a differential selection protocol to uniquely enable identification of context-specific proteolytic events in tissue environments. The PSurf library, comprising over 10^6^ randomized 7-amino-acid sequences, offers 100- to 10,000-fold greater sequence coverage than existing synthetic substrate libraries^27, 32^. We first demonstrated PSurf’s capabilities using recombinant cathepsin B, identifying both canonical and novel substrates with distinct pH sensitivities. We subsequently applied PSurf to murine models of lung metastasis and identified tumor-activated substrates that spared healthy tissues. From these selections, we developed fluorogenic probes exhibiting significantly greater sensitivity and specificity than conventional metalloproteinase-activatable probes. One substrate was further engineered into a membrane insertion probe for *ex vivo* tissue imaging, enabling detection of disseminated lung tumors. *In vivo*, conjugation of top PSurf substrate hits to tumor-trafficking nanobodies enabled multiplexed proteolytic activity assessment via urinary readout, and, when combined with CRISPR–Cas–mediated signal amplification, allowed rapid disease classification distinguishing tumor-bearing from healthy animals.

**Figure 1.**
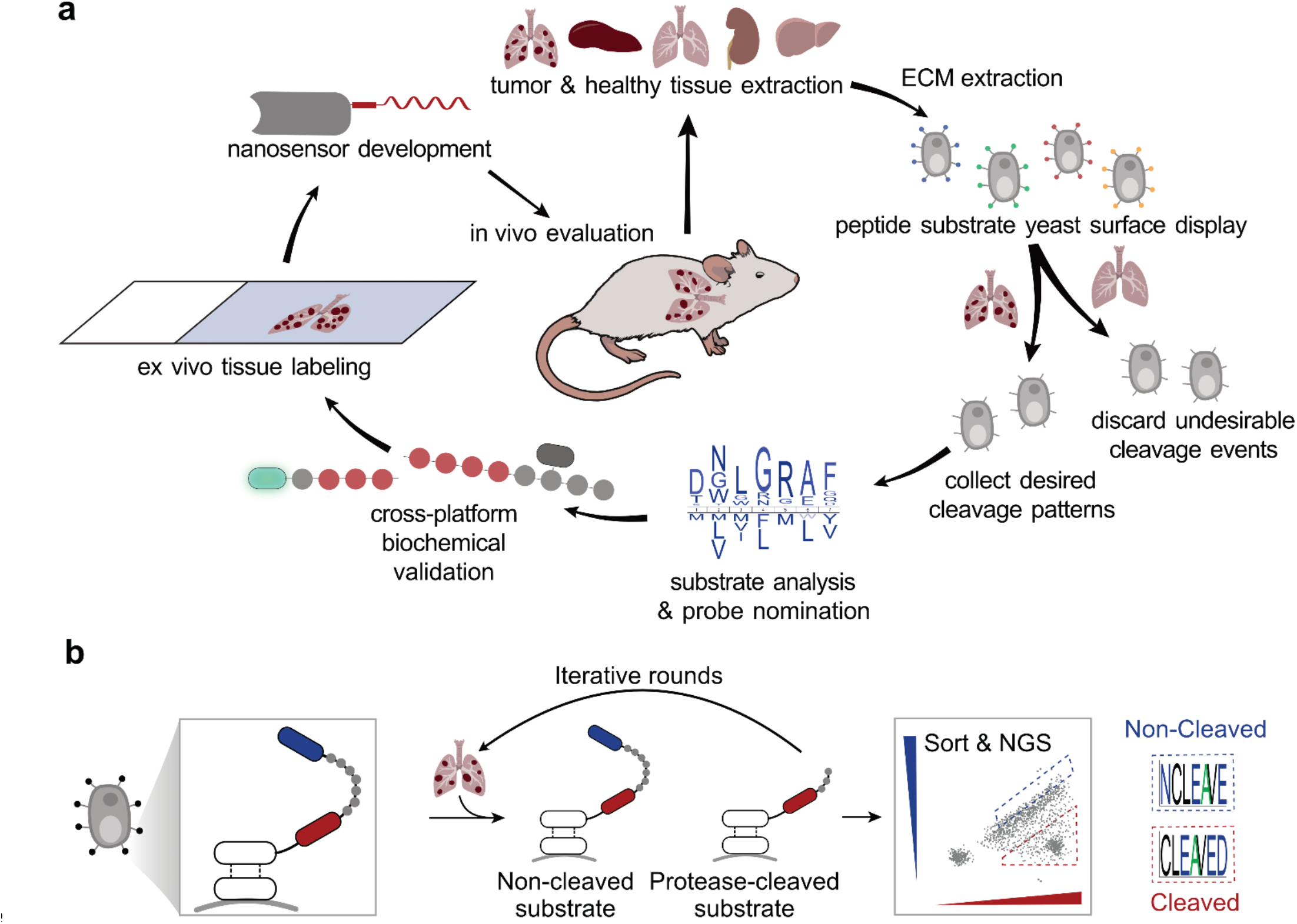
Overview of the PSurf platform for characterizing tissue protease activity. **(a)** Workflow schematic for protease profiling using tissue samples. Organs and tumors are harvested from animal models, and extracellular fluid samples are subjected to substrate profiling using the PSurf yeast display library. Differential selection eliminates non-specific cleavage sequences and enriches tumor-specific substrates. Selected substrates are validated through *in vitro, ex vivo*, and *in vivo* assays, resulting in novel tissue-activated substrate sequences. **(b)** Diagram of the yeast display strategy for tissue protease analysis. Peptides flanked by epitope tags are displayed on the yeast surface. Protease activity removes one tag, and yeast are stained with detection antibodies for analysis and sorting. Customized fluorescence-activated cell sorting (FACS)-based, multi-round selection strategies generate an enriched yeast pool, which is amplified and analyzed by next-generation sequencing (NGS) to identify tissue-sensitive substrates.

Collectively, our findings establish PSurf as a powerful platform for customizable, high-resolution profiling of proteolytic activity and substrate discovery in biological samples. PSurf offers significant potential to advance understanding of extracellular protease biology and to support the development of protease-activated diagnostic and therapeutic tools.

## Results

### Library design and construction

PSurf employs a double-tagged peptide display approach (**Fig. 1b**). Each peptide displayed on the yeast surface is flanked by two epitope tags recognized by fluorescently-labeled antibodies. When proteolysis occurs, one tag is cleaved off, allowing tracking of cleavage events by measuring fluorescence change and performing cell sorting. Yeast populations displaying peptides with specific proteolytic activities are sorted for NGS analysis or cultured for further rounds of selection. To determine the best tagging strategy, we used the Tobacco Etch Virus (TEV) protease substrate, ENLYFQG. The substrate was fused to the C-terminus of Aga2p for yeast surface display. Three tag pairs were compared: HA/cMyc, HA/V5, and V5/HA (N-/C-terminal orientations) (**Fig. 2a**). Yeast cells displaying the TEV substrate peptides were incubated with TEV protease for 24 hours and analyzed using flow cytometry. Among the designs, the V5/HA tag combination exhibited a clear diagonal distribution in untreated samples, as well as a dose- and time-dependent reduction in HA signal upon proteolysis, indicating homogeneous display and accurate detection of cleavage (**Fig. 2a, b**). In contrast, the HA/cMyc tag pair showed non-uniform staining with a broad spread of cMyc signals, while the HA/V5 configuration showed only a small reduction in V5 signal, failing to reliably reflect TEV cleavage activity (**Fig. 2a**). These results identified the V5/HA tag pair as the most optimal design for constructing the library.

**Figure 2.**
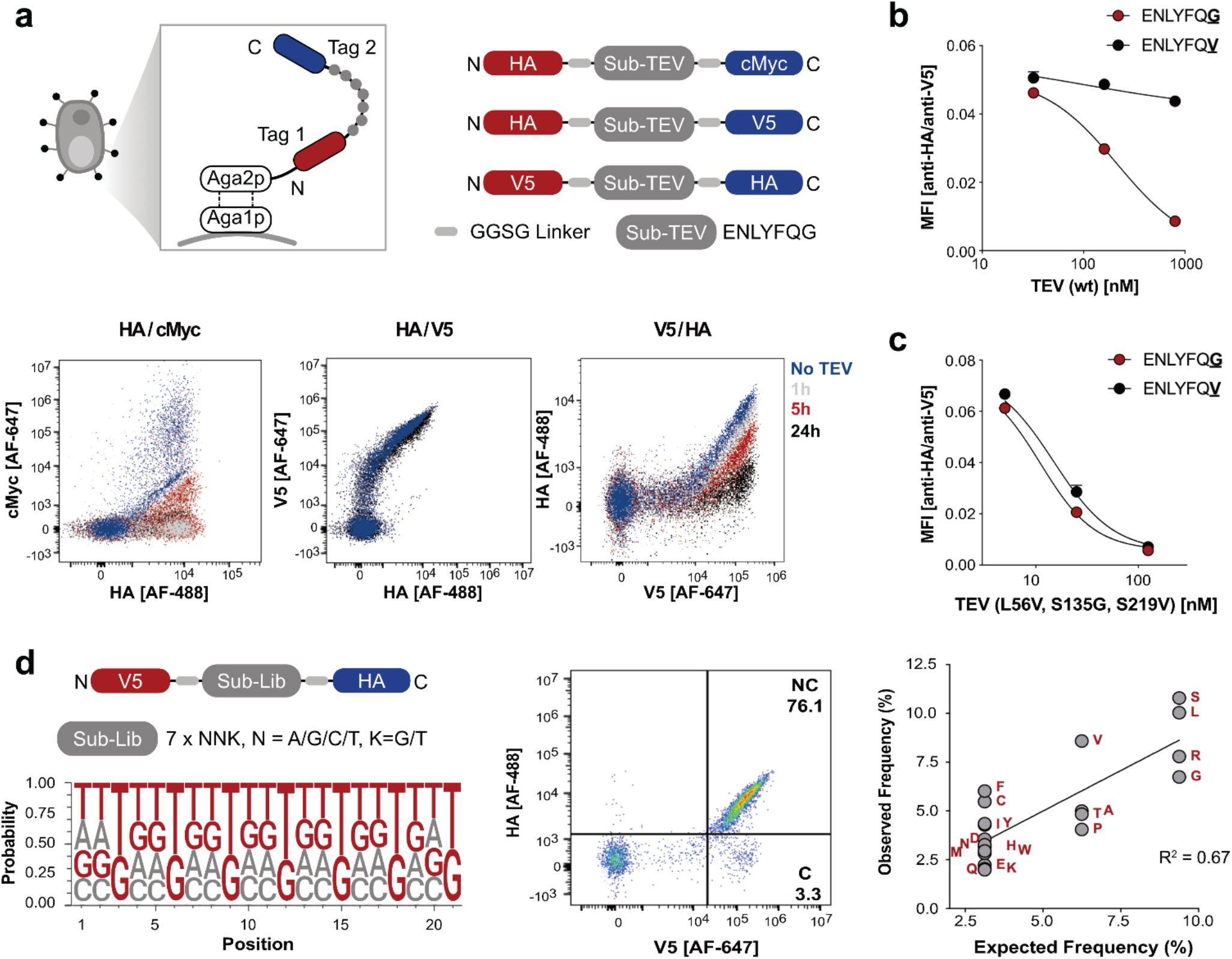
Design, construction, and validation of the yeast library. **(a)** Tag pair selection using a TEV substrate. Schematics and flow cytometry results show three tag pairs flanking a TEV substrate peptide treated with TEV protease over time. The HA/cMyc pair showed uneven cMyc staining but responded to TEV cleavage. The HA/V5 pair displayed uniform staining but no response to protease. The V5/HA pair demonstrated uniform staining and a time-dependent response to TEV protease. **(b)** Validation of the peptide display strategy with TEV substrates. V5- and HA-tag-flanked TEV substrates were incubated with wild-type TEV protease for 24 hours. Yeast display detected sequence-dependent cleavage, with the non-canonical sequence ENLYFQV showing reduced cleavage compared to the canonical ENLYFQG. Data represent the mean of n = 2 replicates; error bars indicate standard deviation. **(c)** A TEV mutant (L56V, S135G, S219V) displayed similar activity toward both canonical and non-canonical substrates. Data represent the mean of n = 2 replicates; error bars indicate standard deviation. **(d)** Design and characterization of the randomized PSurf library flanked by V5 and HA tags. The DNA logo (left) shows base frequencies per position from NGS, consistent with the expected NNK codon distribution. The middle panel indicates 76% of yeast displayed intact peptides with both tags. The right panel shows a strong correlation (R^2^ = 0.67) between observed and expected amino acid distributions for an NNK-based library.

To assess whether the display strategy can differentiate substrates with varying protease sensitivities, we compared the canonical TEV substrate (ENLYFQG) with a non-canonical substrate (ENLYFQV) that has reduced susceptibility to cleavage. Using wild-type TEV protease, we observed significantly reduced cleavage of the non-canonical substrate (**Fig. 2b**). In contrast, cleavage was comparable to the canonical substrate when using an enhanced TEV mutant (TEV L56V, S135G, S219V), consistent with previous studies (**Fig. 2c**). These results validated the ability of the yeast display strategy to detect proteolysis and differentiate substrates of proteases with varying cleavage sensitivities.

A PSurf library displaying randomized seven-amino-acid peptides was then constructed using overlap extension PCR with oligos containing NNK degenerate codons (N: A/T/C/G, K: G/T, **Extended Data Fig. 1a, b**). The peptide length was chosen to balance coverage of central recognition motifs and flanking residues while maintaining sufficient sequence diversity within the practical size constraints of a yeast library^44^. A yeast surface display plasmid pCL2 was linearized and transformed into *Saccharomyces cerevisiae* strain EBY100, along with the amplified DNA library containing homologous flanking sequences to enable recombination. Electroporation generated a library containing 3 × 10^8^ transformants. To characterize the resulting library, DNA from 5x10^6^ transformed yeast cells was amplified and sequenced by NGS. The results showed the nucleotides and amino acid frequencies of the library clones closely aligned with the expected distributions (**Fig. 2d** left panel, **Extended Data Fig. 1d** middle panel). Flow cytometry of the induced library showed that 73% of the yeast population expressed high levels of peptides, with 58% containing both HA and V5 tags (**Extended Data Fig. 1c**).

To understand the cause of the population of yeasts lacking HA, we sorted and sequenced these cells and performed motif analysis. In addition to a fraction of stop-codon–containing peptides resulting from degenerate oligo usage (**Extended Data Fig. 1d**, left panel), 34% of these sequences encoded full-length peptides with strong enrichment of LxxR and RR motifs (**Extended Data Fig. 3b**). These motifs correspond to the known cleavage sequences of the yeast protease *Kex2*, indicating that the peptides were cleaved during secretion to the yeast surface and that our peptide surface display approach unbiasedly identified this enzyme’s substrate motifs (**Extended Data Fig. 3a**)^39^.

The library was further sorted to enrich the non-cleaved population (**Fig. 2d**, middle panel), the relevant subset for tissue proteolysis profiling, and analysis of sequence distribution showed amino acid representation similar to the original library (**Fig. 2d**, right panel; **Extended Data Fig. 1d**, right panel). These results demonstrate successful construction of a diverse yeast surface display peptide library for proteolysis analysis.

### Identifying cathepsin substrates and pH responses

We first applied the library to a recombinant enzyme, human cathepsin B (hCTSB), which is widely used for localized payload delivery and recognized as a prognostic biomarker in multiple solid tumors (**Extended Data Fig. 3a**) ^45, 46^.Cathepsin substrate-based inhibitors show promise as anticancer therapies, while cathepsin-sensitive linkers are commonly employed in antibody-drug conjugates for targeted drug release.^47-49^

pH sensitivity is an important feature of cathepsins, enabling them to modulate activity across cellular compartments and reach peak activity in acidic environments such as lysosomes. hCTSB cleavage was first characterized at pH 4.4. In the first selection round, a 10^7^ yeast culture containing approximately 10^6^ different sequences was treated with 240 nM hCTSB at pH 4.4 for 4h, resulting in 28% cleavage of the displayed peptide. The input library size was chosen based on cell sorting capacity. After removing pre-cleaved peptides from the selected population, a second round of 240 nM hCTSB treatment for 4h led to 63% cleavage (**Fig. 3a, Extended Data Fig. 2a, b** left panel, **Supplementary Fig. S1**). For the third round, the selected population underwent more stringent conditions with reduced hCTSB concentration (122.5, 45, and 10 nM) and shorter incubation time (2 h), yielding a smaller pool of cleaved yeast (**Fig. 3b, Extended Data Fig. 2b**, right panel). NGS analysis revealed strong enrichment of the leucine-valine-glycine (LVG) motif, with the highest enrichment observed under the lowest enzyme concentration. The identification of LVG as a pH 4.4 hCTSB substrate aligned with observations from previous reports^50-53^.

**Figure 3.**
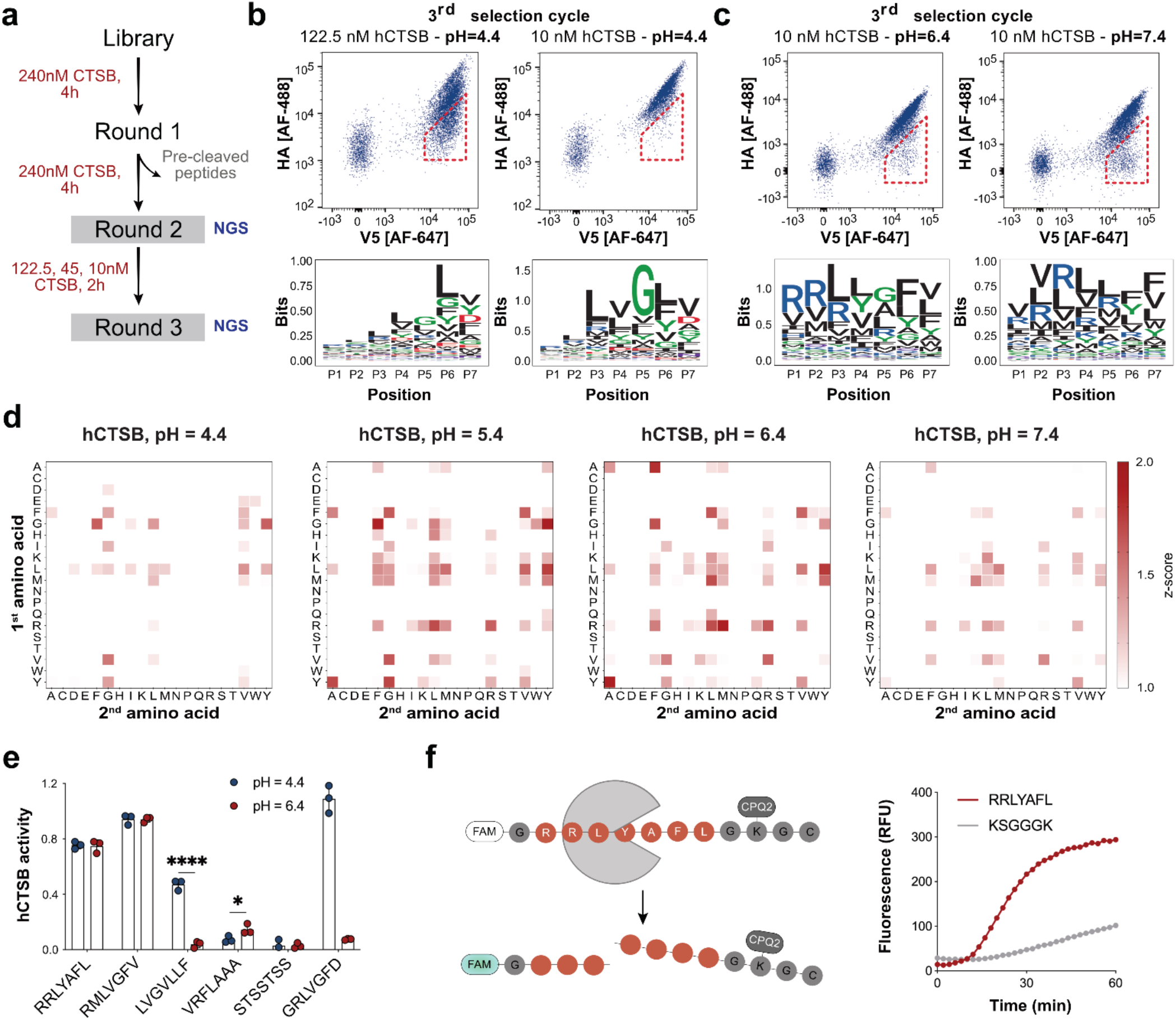
Identifying cathepsin substrates. **(a)** Selection schematic for CTSB substrates identification. **(b)** Flow cytometry plots and sequence logos from PSurf selection with hCTSB at pH 4.4. A strongly enriched LVG cleavage motif was observed under stringent selection conditions (10 nM hCTSB). Sorting gates are indicated by dashed lines. **(c)** Flow cytometry plots and sequence logos showing substrate cleavage motifs at pH 6.4 and 7.4. Unlike the pH 4.4 results, motifs such as RR and VR were strongly enriched. Sorting gates are indicated by dashed lines. **(d)** Z-score plots showing dipeptide enrichment profiles for hCTSB at different pH levels. The y-axis represents the first amino acid; the x-axis the second. Distinct dipeptide motifs were enriched under each pH condition. **(e)** On-yeast cleavage activity of hCTSB with newly identified substrates. RRLYAFL and RMLVGFV were highly sensitive to hCTSB at both pH 4.4 and 6.4; LVGVLLF showed significantly selective activity at pH 4.4, and VRFLAAA at pH 6.4. Yeast displaying these peptides were incubated with 70 nM hCTSB for 72 hours. Cleavage activity is shown as the negative log of HA-AF488 signal normalized to peptide expression (V5-AF647 signal) and to untreated controls. Data represents the mean of n = 3 replicates; error bars indicate standard deviation. STSSTSS served as a no-activity control and GRLVGFD^53^ served as a positive control. Unpaired *t*-test was used for pH-dependent activity comparisons and p-values are presented in Table S4. **(f)** Schematic of the probe and cleavage assay comparing RRLYAFL to the conventional substrate KSGGGK. A synthetic peptide labeled with FAM/CPQ2 was used; cleavage releases the fluorophore, generating a fluorescence signal. RRLYAFL showed enhanced cleavage by hCTSB. Data represents the mean of n = 3 replicates.

We then investigated hCTSB’s pH-dependent substrate preferences by performing similar selection cycles at pH 5.4, 6.4, and 7.4 (**Fig. 3c, Extended Data Fig. 2c**). Compared to the pH 4.4 selections, smaller percentages of substrate cleavage were observed at higher pH values, reflecting the expected decrease in hCTSB activity at more neutral pH^54^. NGS results indicated that as pH increased, enrichment of the LVG motif decreased, while RR and VR motifs became more prominent (**Fig. 3d, Extended Data Fig. 2d, e**). RR and VR has previously been identified as a sensitive hCTSB substrate at pH 7.0 and 6.4, in line with our findings^55, 56^. Notably, a VR-derivative peptide, Valine-Citrulline, is one of the most widely used linkers in antibody-drug conjugates^57^. These results highlight the ability of PSurf to identify context-specific enzyme cleavage features.

Since cathepsins mainly promote tumor progression in the extracellular space, where pH is typically above 5.5, there is strong interest in identifying novel cathepsin substrates or peptide-based inhibitors that function at higher pH to better target tumor-associated cathepsins. We therefore cloned four peptides, either directly identified from the pH 6.4 screen or designed by combining top enriched motifs, and tested their cleavage activity over time on yeast (**Fig. 3e, Table S2**). Consistent with the motif enrichment results, the VR-containing peptide VRFLAAA showed selectivity at pH 6.4, while the LVG-containing peptide LVGVLLF showed selectivity at pH 4.4 (**Fig. 3e**). Peptide RMLVGFV, containing the RM motif enriched at pH 6.4 and the LVG motif enriched at pH 4.4, was active at both pH values. Peptide RRLYAFL, containing the RLY motif enriched at both pH levels (**Fig. 3b, c**), also showed activity at both pH conditions.

From our analysis, the novel substrate RRLYAFL emerged as the most enriched sequence in the pH 6.4 NGS dataset, representing 12% of the NGS sequences. To assess its activity relative to known cathepsin substrates, we designed synthetic Förster resonance energy transfer (FRET)-paired substrates, consisting of 7 amino acids flanked by a fluorophore and quencher, where cleavage leads to time-dependent fluorescence increase (**Fig. 3f**, left panel). Testing this probe alongside a 6-amino acid CTSB-responsive probe, we found that the newly identified substrate exhibited faster cleavage kinetics and greater fluorescence signal than the known CTSB substrate at a pathological extracellular pH (6.4) (**Fig. 3f**, right panel). These results demonstrate that our PSurf library can effectively identify both known and novel cathepsin substrates.

Samples derived from living organisms exhibit greater complexity than purified proteases due to the presence of diverse protease families, endogenous inhibitors, competing substrates, metal ions, and other components that modulate protease activity. To evaluate the PSurf system’s ability to characterize such complex samples, we applied it to a mouse upper intestinal washout, which contains high concentrations of trypsins expected to cleave Arg-rich sequences (**Extended Data Fig. 3a, c; Supplementary Fig. S2**). Incubation of the PSurf library with this sample for 24 hours resulted in cleavage of 24% of displayed peptides, with NGS analysis revealing significant arginine enrichment in the cleaved pool, in contrast to the non-cleaved fraction (**Extended Data Fig. 3d**). A subsequent selection cycle with a reduced 4-hour incubation yielded 10% cleavage and further arginine enrichment across all seven amino acid positions (**Extended Data Fig. 3e; Supplementary Fig. S2**). These results demonstrate that PSurf can effectively capture specific proteolytic activities in biological samples and that increased selection stringency enhances enrichment for tissue-sensitive substrates, consistent with observations in the hCTSB experiments.

### Identify tumor-selective sequences through differential selection

We next applied PSurf to tumor samples. We harvested tumor bearing lungs and healthy organs from two lung metastasis mouse models: metastatic colorectal cancer (based on CT26 colon adenocarcinoma cells) and metastatic melanoma (based on B16-F10 melanoma cells). In these models, tumor cells were injected into the tail vein, leading to lung colonization and visible metastatic nodules within 10–14 days. We extracted the soluble fraction of the ECM from healthy tissues (liver, lung, kidney, and spleen) and tumor lesions in the lung based on a filter centrifugation method previously applied in metabolomics studies (**Fig. 4a, Extended Data Fig. 4a, see Methods**)^58^. Alternatively, in some contexts, we prepared tissue homogenates by mechanical dissociation and collected extracellular proteins for downstream assays.

**Figure 4.**
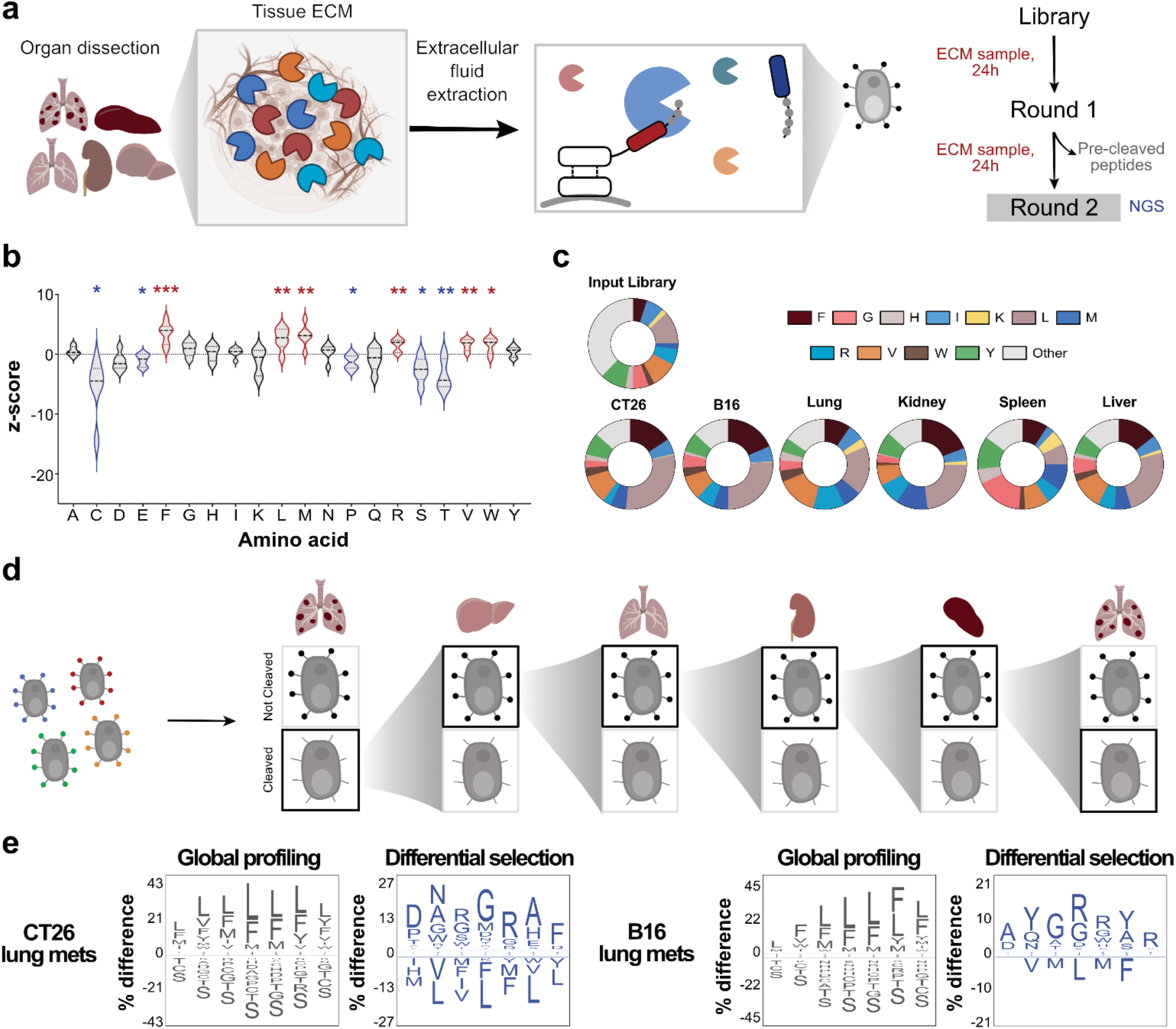
Profiling of tissue extracellular fluids using PSurf. **(a)** Schematic and protocol overview for isolating and profiling tissue-derived extracellular fluids. **(b)** Z-score plots showing single amino acid enrichment and depletion across tissues and tumors (liver, lung, kidney, spleen, CT26 CRC lung metastases, and B16 lung metastases). Residues with a significant increase compared to pooled mean of Z-scores are framed in red, indicating enrichment. Residues with a significant decrease compared to the pooled mean are framed in blue, indicating depletion. p-values are provided in Table S4. **(c)** Frequencies of enriched amino acids in the input library and each sample type. **(d)** Schematic of the differential selection strategy. The tumor-enriched population was sequentially screened against liver, lung, kidney, and spleen samples. After each round, the non-cleaved pool was carried forward to remove substrates cleaved by healthy tissues. The final non-cleaved pool was screened against tumor extracellular fluid to further enrich for tumor-sensitive substrates. **(e)** IceLogos showing per-position amino acid enrichment and depletion before and after differential selection, normalized to the initial library. The pool selected from global profiling was enriched in leucine and phenylalanine, while the final pool following differential selection displayed greater sequence diversity. IceLogos were generated using the IceLogo web server.

The PSurf library was incubated with ECM samples for 24 hours, resulting in substrate cleavage levels of 11% in CT26 tumor-bearing lung samples, 10% in B16 tumor-bearing lung samples, and 7% in healthy lung samples (**Fig. 4a, Extended Data Fig. 4b, Supplementary Fig. S3**). Healthy spleen, kidney, and liver exhibited cleavage of 8%, 9%, and 10%, respectively. The buffer control led to 6% cleavage, reflecting residual pre-cleaved peptides by yeast, though the mean fluorescence intensity indicated a reduced degree of cleavage compared to tissue-derived samples. The cleaved fractions were harvested, pre-cleaved peptides were removed by sorting, and the resulting yeast pool was then subjected to a second round of selection with tissue ECM samples and analyzed by NGS sequencing.

Interestingly, the NGS results revealed strong enrichment of sequences containing phenylalanine (Phe), leucine (Leu), and other hydrophobic residues across both healthy tissue and tumor samples (**Fig. 4b, c; Extended Data Fig. 4c, d**). Such Phe- and Leu-rich motifs are characteristic of substrates for several serine proteases^59^. For example, granzyme H and cathepsin G possess chymotrypsin-like activity and preferentially cleave after large hydrophobic residues like Phe, as evidenced by their efficient cleavage of the Phe-Leu-Phe synthetic substrate^60, 61^.

Cathepsin G further cleaves substrates such as Ala-Ala-Pro-Leu and Ala-Ala-Pro-Phe, sequences with Leu and Phe at the P1 position^62^. Beyond these enzymes, other secreted serine proteases, such as kallikrein-related peptidase 7 (KLK7)^63^, neutrophil elastase (NE1), and proteinase 3 (PR3)^64^, also exhibit substrate preferences for Leu or Phe. In addition, some MMPs (MMP1-3, MMP7-9, MMP12-14) have been reported to prefer leucine at the P1′ position^20^. Thus, the observed enrichment of Phe/Leu-rich sequences may reflect the integrated activity of certain protease classes, such as serine proteases and MMPs, acting within the ECM of healthy and tumor tissues. These findings suggest that tissue samples may share common cleavage signatures shaped by conserved protease substrate preferences. Further investigation into the mechanisms underlying these sequence enrichments could yield important insights into the proteolytic landscapes of normal and pathological tissue environments.

To distinguish tumor-specific substrates from those commonly cleaved in both tumor and healthy tissues, we leveraged yeast display’s capacity for repeated positive and negative selection, a strategy used in nanobody and antibody discovery to isolate binders targeting specific epitopes or conformational states of antigens^65-67^ (**Fig. 4d**). We initiated this differential selection process using the cleaved population obtained from the initial tumor ECM screens (**Extended Data Fig. 5a, left panel**). To deplete substrates also cleaved by healthy ECM, this pool was sequentially incubated with ECM from healthy liver, lung, kidney, and spleen, collecting the non-cleaved fraction at each step and culturing for the next selection step. Finally, the remaining non-cleaved pool was exposed to tumor-derived ECM, and the resulting cleaved fraction was collected, enriching for substrates highly sensitive and specific to tumor-associated proteolytic activity (**Extended Data Fig. 5a, right panel; Supplementary Fig. S4**).

NGS was performed on the non-cleaved and cleaved populations from each selection round. IceLogos^68^ generated from each pool revealed distinct tissue-specific cleavage features (**Extended Data Fig. 5b**). Comparison of IceLogos from the cleaved populations before (input) and after (output) differential selection showed clear differences (**Fig. 4e**). While Leu and Phe were highly enriched in the input sequences from both tumor models, these residues were depleted in the output population. In the final cleaved pool from tumor samples, the proportion of sequences containing Phe, Leu, and related dipeptides was significantly reduced. Instead, residues such as arginine, proline, asparagine, glycine, alanine, and aspartate were enriched, along with new motifs (**Extended Data Fig. 6**).

We selected 10 substrate sequences (**Supplementary Table S3**) and assessed their tumor versus healthy lung cleavage as individually displayed peptides on yeast. Based on the results, four peptides, DNLGRAF, PARMLHI, AYGRLYR, and PFLYLFG, exhibited highest tumor-preferred cleavage activity and were selected for further biochemical and *in vivo* analyses (**Extended Data Fig. 5c**).

### Biochemical characterization of PSurf-identified substrates

To further characterize the PSurf-identified sequences, we synthesized these peptides as fluorogenic FRET probes. Lung homogenates from mice bearing CT26 metastases and healthy control animals were incubated with the probes to assess differences in substrate cleavage rates (**Fig. 5a**). Inflamed lung samples from lipopolysaccharide-treated mice were also included for comparison to distinguish tumor-specific from inflammation-associated proteolytic activity. The four substrates identified by yeast display were benchmarked against the previously reported MMP substrates PVPLSLVM and PLGLRSW^69, 70^, which contain the conserved MMP cleavage motif PXXL^71^, as well as AIEFSD, a known substrate of granzyme B^72^. The kinetic fluorescence assays revealed that all four PSurf-identified probes were cleaved more rapidly in tumor samples than in healthy lung samples (**Fig. 5b, c**). In particular, the substrates DNLGRAF and AYGRLYR exhibited markedly enhanced tumor cleavage kinetics compared to both the MMP substrates and the granzyme B substrate, achieving superior discrimination between tumor and healthy tissues. Notably, all four substrates exhibited minimal cleavage in inflamed lung tissue, demonstrating their tumor selectivity (**Fig. 5c**).

**Figure 5.**
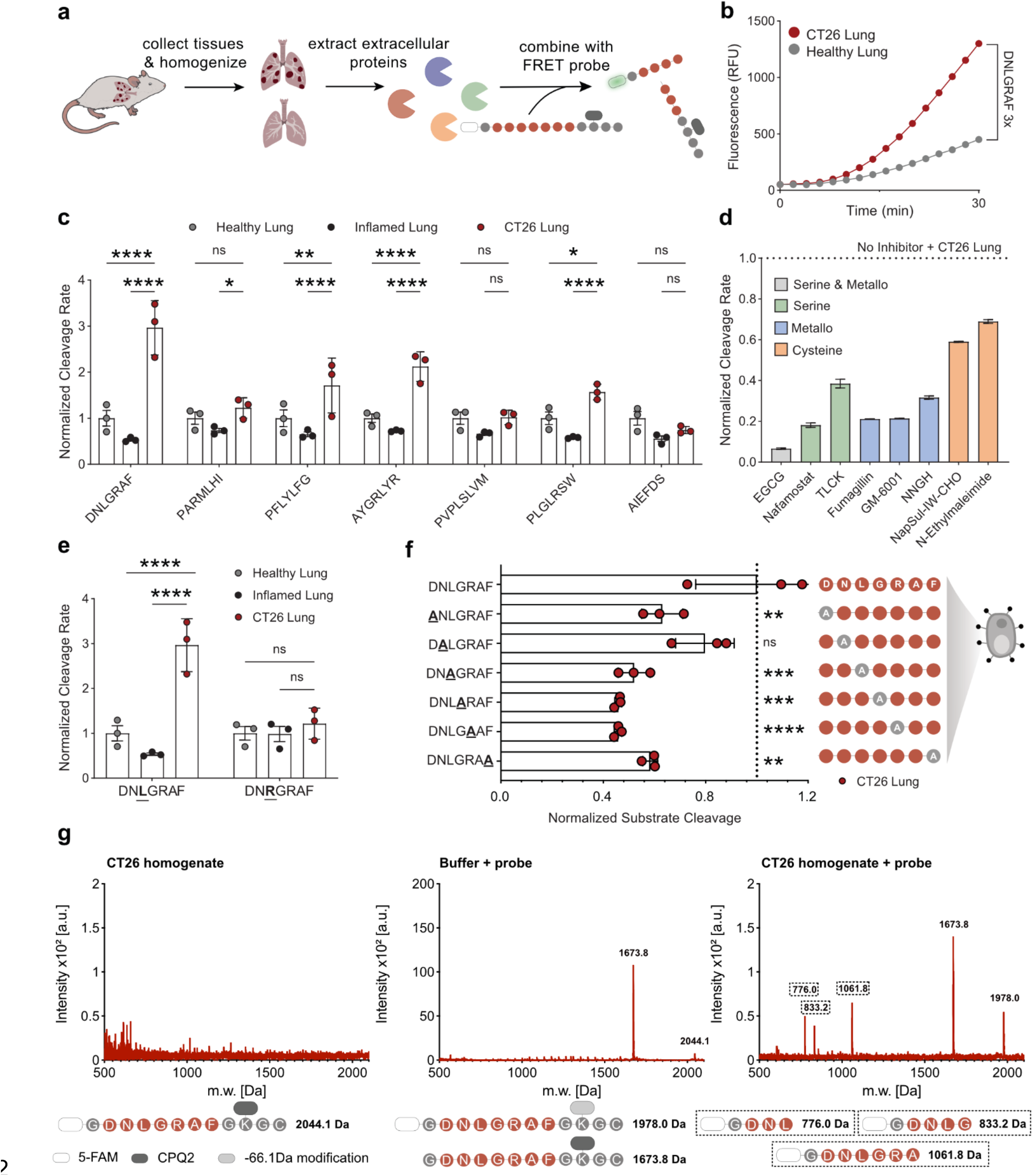
Substrates identified from PSurf exhibit tumor specificity and differ from canonical MMP substrates. **(a)** Schematic illustrating the use of fluorogenic probes to assess substrate cleavage by tissue homogenates. **(b)** Representative kinetic curves showing increasing fluorescence signal from cleaved probes over time. Substrate DNLGRAF demonstrates enhanced activity in response to tumor-bearing lung compared to healthy lung tissue. Data represents mean from n = 3 replicates. **(c)** Bar graphs comparing cleavage kinetics of four PSurf-nominated probes and two known MMP probes and a GZMB probe, respectively in CT26 tumor-bearing, inflamed, and healthy lung homogenates. The slope of the linear region of each kinetic curve was used to calculate cleavage rates across samples. Data represents the mean ± s.e.m. of n = 3 replicates. Two-way ANOVA with Dunnett’s multiple comparisons test was performed and show ***P < 0.0001 for DNLGRAF in CT26 versus healthy lung or inflamed lung (see Table S4). **(d)** Proteolysis inhibition assay compares DNLGRAF cleavage in the presence of inhibitors (150 µM) known to inhibit different protease families. Data represents the mean ± s.e.m. of n = 3 replicates. **(e)** Bar graphs comparing cleavage kinetics of logo-derived peptide (DN<u>R</u>GRAF) with the most enriched peptide (DN<u>L</u>GRAF), showing the better performance of the latter. DNLGRAF data are the same as in panel c and are included again for comparison. Data represents the mean ± s.e.m. of n = 3 replicates. **(f)** Bar graphs comparing on-yeast cleavage activity of positional alanine scanning mutants of DNLGRAF. Variants with each position individually replaced by alanine were treated with CT26 lung homogenates for 24 h. Cleavage signals were normalized to the signal of the intact DNLGRAF substrate. Data represent the mean ± s.d. of n = 3 replicates. One-way ANOVA with multiple comparisons was performed and significance was defined as P < 0.05 (see Table S4). **(g)** Mass spectrometry analysis of CT26 homogenate alone (left), DNLGRAF probe in assay buffer (middle), and DNLGRAF probe treated with CT26 homogenate (right). The CT26-treated DNLGRAF probe shows three unique peaks corresponding to cleaved probe fragments (highlighted with dashed frames). Schematic illustrations of the probe fragments and their corresponding observed mass peaks are shown at the bottom.

We next sought to identify which tumor-associated protease families contribute to the substrate cleavage. Analysis of transcriptomic data from OncoDB^73^ revealed upregulation of proteases from multiple families, including cysteine, metallo, and serine proteases, in CRC samples compared to healthy colon tissue (**Supplementary Fig. S5a**). We first assessed the cleavage specificity of the four newly identified substrates, alongside the canonical MMP substrate PVPLSLVM, using recombinant catalytic domains of MMPs 1–3 and 7–14 (**Extended Data Fig. 7a**). As expected, PVPLSLVM was cleaved by various MMPs. In contrast, three of the PSurf-identified substrates, DNLGRAF, PFLYLFG, and AYGRLYR, were not cleaved by any of the tested MMPs. Additionally, these substrates were not cleaved by serine proteases upregulated in CRC, including FAP^17^, GZMB^72^, and PCSK9^74^, nor by the cysteine protease CTSB, enzymes that have frequently been targeted in recombinant enzyme-based substrate discovery. Among the four new substrates, PARMLHI was the only sequence partially cleaved by certain MMPs (**Extended Data Fig. 7a**).

We further studied the related protease families using protease inhibitors (**Supplementary Table S6**). The inhibitor panel did not reveal exclusive sensitivity of the substrates to a single protease or protease class; rather, cleavage was most significantly reduced by inhibitors targeting serine and metalloprotease families in combination (**Fig. 5d, Extended Data Fig. 7b, Supplementary Fig. S5b**). Specifically, cleavage of DNLGRAF and AYGRLYR was strongly inhibited by the broad-spectrum serine and metalloprotease inhibitor epigallocatechin gallate (EGCG) and by the serine protease inhibitor nafamostat, while showing minimal response to cysteine protease inhibitors. PARMLHI, consistent with its partial cleavage by MMPs, was inhibited by MMP inhibitors such as marimastat and phosphoramidon. In contrast, PFLYLFG was more strongly inhibited by cysteine protease inhibitors, indicating involvement of proprotein convertases and suggesting distinct protease sensitivity compared to the other substrates. These findings indicate that tissue selection identified substrates are responsive to multiple tumor-associated proteases and represent distinct classes of protease-reactive probes compared to conventional MMP substrates.

To investigate how amino acid composition affects substrate sensitivity, we focused on the PSurf-identified substrate with highest tumor sensitivity, DNLGRAF. We first synthesized a fluorogenic probe with the sequence DNRGRAF. This sequence was designed by selecting at each position the most enriched amino acid from the differential selection NGS data (as shown in the IceLogo of **Fig. 4e**). Notably, DNRGRAF differs from the DNLGRAF sequence by only a single residue at the P3 position (R to L). However, when tested against CT26 lung homogenates, DNRGRAF displayed substantially lower activity compared to DNLGRAF (**Fig. 5e**). This result revealed the importance of position P3 for efficient cleavage of this substrate.

To further investigate the contribution of individual amino acid positions, we performed an alanine scan on the DNLGRAF substrate. Each of the six non-alanine residues at positions 1–5 and 7 was individually substituted with alanine, and cleavage activity was assessed using yeast display in the presence of CT26 homogenate samples (**Fig. 5f**). All alanine variants exhibited reduced cleavage compared to the original DNLGRAF sequence, indicating that all positions contribute to protease recognition or cleavage activity. Particularly, substitutions at positions 3, 4, and 5 (LGR) resulted in the most pronounced decreases in substrate sensitivity.

Lastly, we used matrix-assisted laser desorption ionization–time of flight mass spectrometry (MALDI-TOF MS) to identify specific cleavage sites within DNLGRAF. CT26 homogenate was incubated with the fluorogenic DNLGRAF probe for one hour, alongside controls that either lacked the probe or contained the probe alone treated under identical conditions. As expected, CT26 homogenate alone showed no probe-derived peaks above 500 Da (**Fig. 5g, Extended Data Fig. 7c**). The probe alone displayed peaks at 2044.1 Da (full-length probe) and 1673.8 Da (w/o 5-FAM fluorophore) (**Fig. 5g**). After incubation with CT26 homogenate, the DNLGRAF probe exhibited the same 1673.8 Da peak seen in the untreated probe but also produced four additional significant peaks at 776.0, 833.2, 1061.8, and 1978.0 Da, while the 2044.1 Da peak corresponding to the intact probe disappeared (**Fig. 5g**). Three of these peaks, 776.0 Da (5FAM-G-D-N-L; expected m/z = 777.2), 833.2 Da (5FAM-G-D-N-L-G; expected m/z = 834.2), and 1061.8 Da (5FAM-G-D-N-L-G-R-A; expected m/z = 1061.4), correspond to fragments consistent with cleavage sites between positions 3/4, 4/5, and 6/7, respectively (**Fig. 5e**). The peak at 1978.0 Da represents a loss of ∼66.1 Da from the intact probe mass (2044.1 Da), potentially due to cleavage within the CPQ2 quencher moiety. To confirm that these fragments arose from tissue-derived activity, we monitored the accumulation of higher-mass fragments (>1000 Da) over time. Peaks at 1061.8 and 1978.0 Da increased in intensity during incubation, supporting time-dependent probe cleavage and modification by CT26-derived enzymes (**Extended Data Fig. 7c, d**). These results revealed specific cleavage sites within the DNLGRAF substrate.

Collectively, these biochemical characterizations demonstrate that PSurf-identified substrates are selectively cleaved by proteases active in tumor tissue and that protease recognition and cleavage depends on the precise sequence context across the substrate. Furthermore, the finding that a substrate designed from position-by-position enrichment exhibited lower activity than DNLGRAF indicates that substrate nomination may benefit from using sequences identified through selection rather than relying solely on motif-based design.

### Tumor-activated probe identifies metastases with spatial resolution

To determine spatial proteolytic activity in intact tissue slices, we designed a probe based on the DNLGRAF substrate that enables both fluorescence activation and local membrane insertion upon cleavage. Specifically, DNLGRAF was fused to a membrane-binding peptide (FVQWFSKFLLG)^75^, with a Cy7 fluorophore at the N-terminus and a QC1 quencher at the C-terminus, enabling cleavage-dependent fluorescence activation. In the intact probe, the proximity of Cy7 and QC1 results in fluorescence quenching. Cleavage by tissue proteases separates the fluorophore from the quencher, leading to dequenching and restoration of Cy7 fluorescence. Simultaneously, the liberated probe inserts into cell membranes, allowing spatial localization of protease activity (**Fig. 6a**).

**Figure 6.**
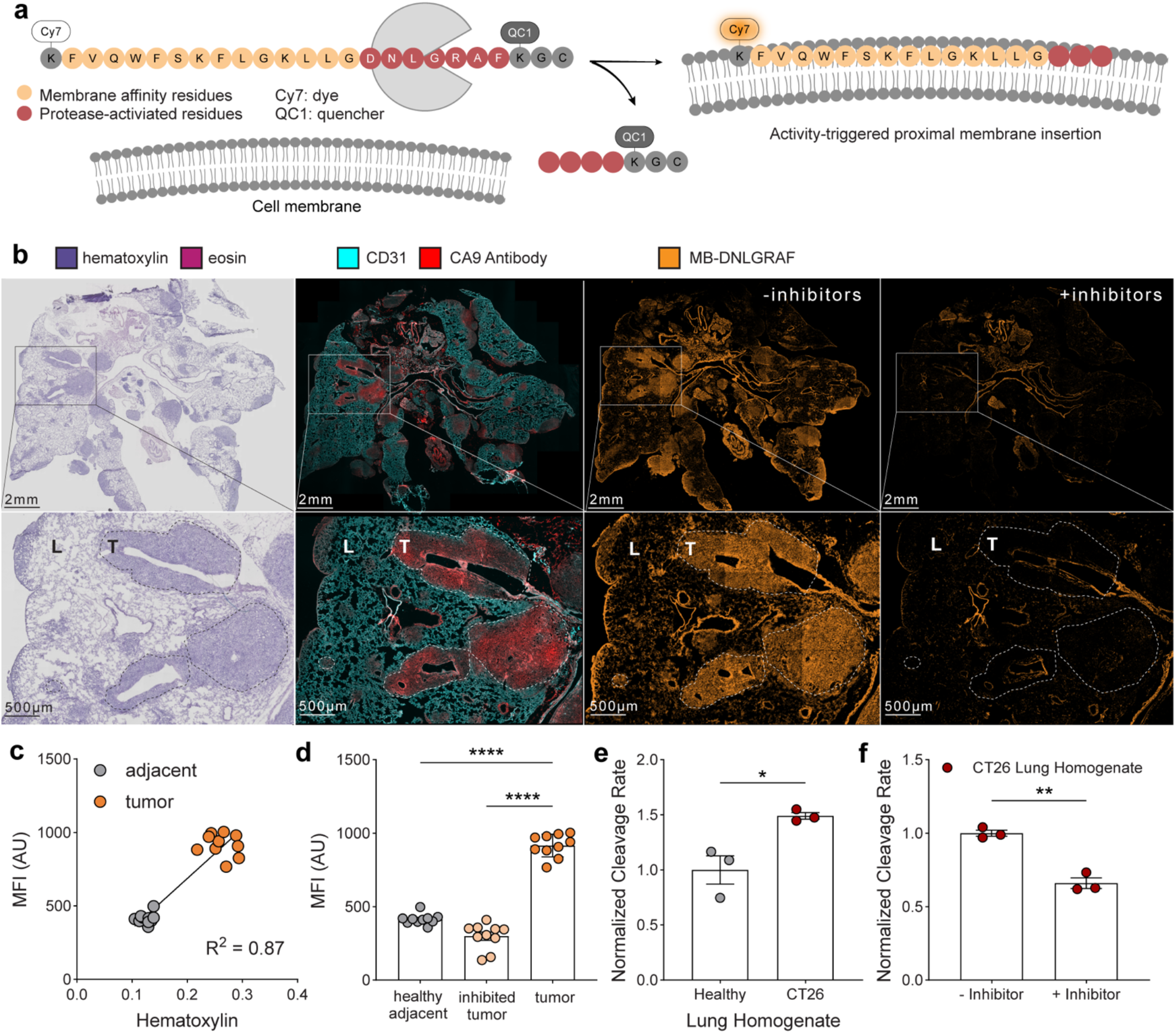
New substrate-based spatial probe delineates CT26 metastases in whole tissue sections. **(a)** Schematic of MB-DNLGRAF, a membrane-binding fluorogenic probe that emits fluorescence following proteolytic cleavage of the DNLGRAF substrate. In the intact probe, Cy7 fluorescence is quenched by QC1; cleavage separates the fluorophore and quencher, enabling membrane insertion and fluorescence activation. **(b)** Representative images of mouse lung tissue with CT26 metastases stained with hematoxylin and eosin, anti-CA9 or anti-CD31 antibodies, or the MB-DNLGRAF probe with or without a protease inhibitor cocktail. Hematoxylin (blue), eosin (red), CA9 (red), CD31 (vasculature, cyan), MB-DNLGRAF (orange). L: healthy lung; T: tumor nodule. **(c)** Correlation analysis of hematoxylin and MB-DNLGRAF fluorescence signals shows a strong positive correlation, indicating accurate tumor delineation by the MB-DNLGRAF probe. Signals were quantified from ROIs drawn across lung sections (see **Supplementary Fig. S6**). **(d)** Bar graphs showing mean fluorescence intensity (MFI) of MB-DNLGRAF in tumor lesions compared to healthy tissues. Signal is significantly higher in tumor lesions compared to adjacent healthy tissue. Data represent mean ± s.e.m. across n = 10 ROIs; two-way ANOVA with Tukey’s multiple comparisons test, ****P < 0.0001 for tumor versus healthy adjacent tissue, ****P < 0.0001 for tumor versus inhibitor-treated tumor. **(e)** Bar graph showing MB-DNLGRAF cleavage rates in solution using healthy lung and CT26 lung homogenates. Data represent mean ± s.e.m. of n = 3 lungs per group; unpaired two-tailed *t*-test, *P = 0.0205 for healthy versus CT26. (**f**) Bar graph showing MB-DNLGRAF cleavage rates in solution using CT26 lung homogenates, with and without protease inhibitors. Probe activation is significantly higher in tumor homogenates and is reduced by 300 µM inhibitor GM6001. Data represent mean ± s.e.m. of n = 3 lungs per group; unpaired two-tailed *t*-test, **P < 0.0012 for CT26 without inhibitor versus CT26 inhibitor-treated.

We applied this membrane-binding DNLGRAF fluorogenic probe, termed MB-DNLGRAF, to stain CT26 lung metastasis tissue sections (**Fig. 6b**). Hematoxylin and eosin staining identified metastatic lesions, and regions of interest (ROIs) were used to correlate hematoxylin signal with probe fluorescence. MB-DNLGRAF fluorescence showed a strong positive linear correlation with hematoxylin staining (R^2^ = 0.87) (**Fig. 6c, Supplementary Fig. S6**). Mean fluorescence intensity was 2.2-fold higher in tumor lesions compared to healthy lung regions (**Fig. 6d**). To confirm protease-dependent probe activation, we applied a cocktail of inhibitors identified from the protease screen to tissue sections, which reduced tumor-associated fluorescence by 67.3% (**Fig. 6d**). This result demonstrates that probe activation in the tumor is protease-dependent.

This probe format was also validated in solution using homogenates from healthy lungs and CT26 metastasis-bearing lungs. Consistently, cleavage rates were significantly higher in tumor homogenates compared to healthy tissue, and activity was markedly reduced in the presence of inhibitors (**Fig. 6e-f**). Together, these results demonstrate that the PSurf-identified substrate DNLGRAF undergoes tumor-specific protease cleavage both in solution and in tissue, enabling spatial detection of proteolysis activities.

### *In vivo* protease–activated biosensors enable disease classification via urinary readout

We next leveraged the tumor-activated protease substrates we identified to engineer biosensors for noninvasive tumor detection. To make the biosensor panel, the four PSurf-identified substrates were fused to a tumor-specific nanobody through a rigid linker to minimize steric hindrance to protease access. A single-stranded, chemically stabilized DNA barcode was conjugated to the biosensor to enable multiplexing and to convert protease activity into a measurable and amplifiable signal (**Fig. 7a**). Following intravenous administration, the biosensor accumulates in the TME through nanobody-mediated targeting. Tumor-associated proteases would cleave the substrate, releasing the DNA barcode, which re-enters circulation, is filtered by the kidneys, and concentrated in the urine, where it can then be noninvasively detected through a CRISPR-Cas12a detection method as an indicator of tumor presence (**Fig. 7b**)^69^.

**Figure 7.**
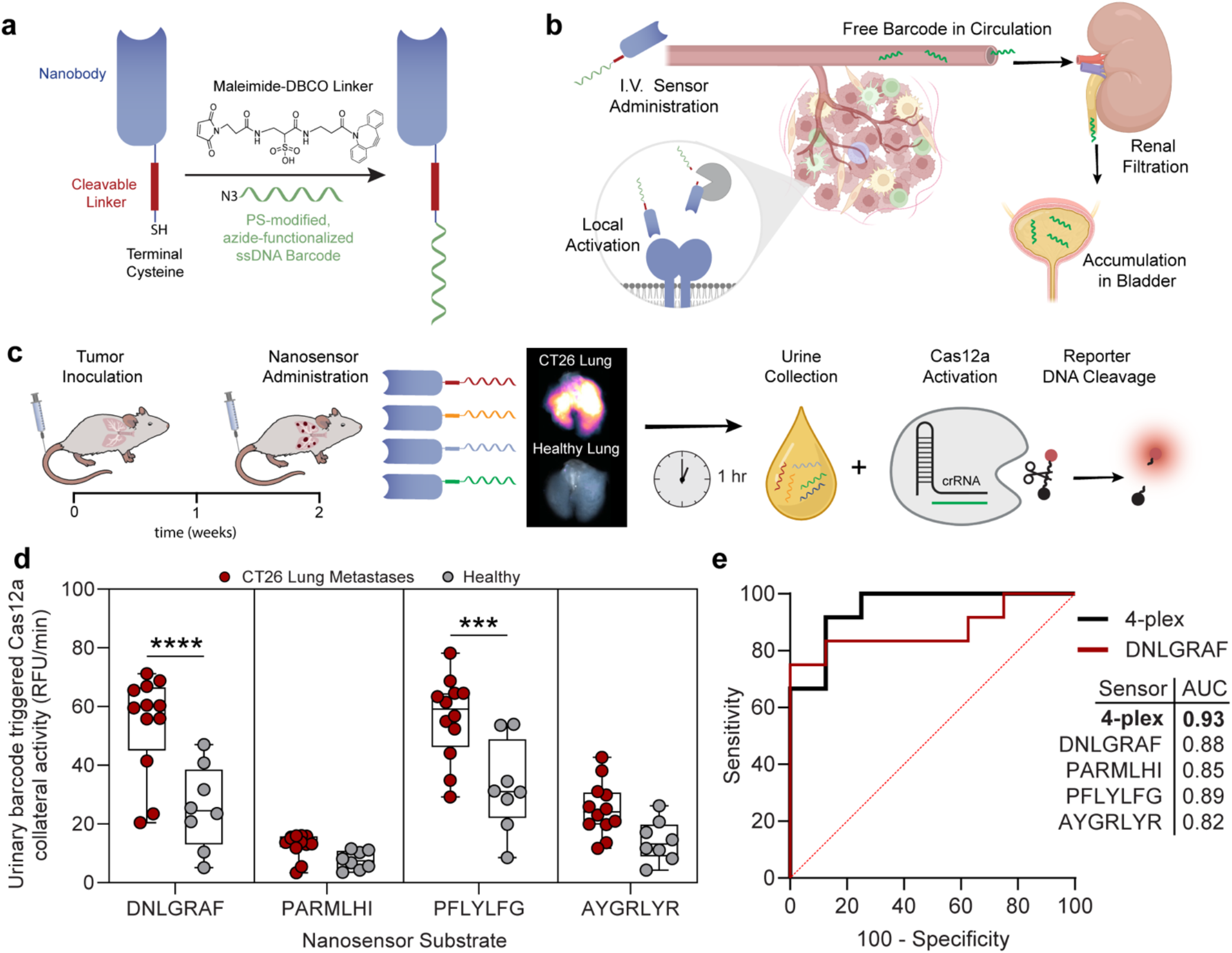
*In vivo* activation of tumor-targeted PSurf substrates reports disease state via urinary readout. **(a–c)** Schematics of the nanosensor design and experimental workflow. **(a)** The nanosensor consists of a CA9-targeting nanobody fused to a protease-cleavable substrate conjugated to a Maleimide-DBCO linker and a DNA barcode for detection. **(b)** Following intravenous injection, nanosensors accumulate in tumor nodules via nanobody targeting. Tumor-associated proteases cleave the substrate linker, releasing the DNA barcode into circulation. Free barcodes are filtered by the kidneys and concentrated in the urine. **(c)** Mice with and without CT26 lung metastases were administered a cocktail of four nanosensors incorporating the DNLGRAF, PARMLHI, PFLYLFG, and AYGRLYR substrates. Representative images of nanobody lung accumulation. Urine was collected over 1 hour post-injection and analyzed for barcode concentration using Cas12a activation and a fluorogenic reporter assay. **(d)** Box plot comparing Cas12a activation, measured by reporter DNA cleavage rate, for each corresponding nanosensor substrate. n = 12 mice per tumor group; n = 8 mice per healthy group; one-way ANOVA with Sidak’s multiple comparisons test: DNLGRAF, ****P < 0.0001; PFLYLFG, ***P = 0.0002. Box plots display the interquartile range from the first quartile (Q1, 25th percentile) to the third quartile (Q3, 75th percentile), with the median (50th percentile) represented by a line within the box. Whiskers extend to the minimum and maximum values. **(e)** ROC analysis demonstrating the nanosensor panel’s ability to classify disease status using a 4-plex multiple linear regression model compared to DNLGRAF alone. The dashed red line indicates a random classifier with an AUC of 0.5.

For tumor targeting, we used a nanobody against carbonic anhydrase IX (CA9). CA9 is one of the most overexpressed antigens in CRC and other solid tumors, playing a key role in mediating tumor acidification (**Extended Data Fig. 8a**). It was selected as our target antigen due to its high expression and the correlation between tumor acidification and the alteration of protease activity^76-80^. To validate CA9 expression, we analyzed transcriptomic data across nine cancer types with high rates of lung metastasis, as well as two types of primary lung cancer. In colon adenocarcinoma (COAD), CA9 was upregulated 53-fold in primary tumors compared to healthy tissue (85.0 ± 163.0 vs 1.6 ± 1.5; P = 0.0012, unpaired two-tailed *t*-test) (**Extended Data Fig. 8b**). This upregulation was consistent across disease stages. In CT26 murine lung metastasis model, histological staining with an anti-CA9 antibody confirmed high CA9 expression in tumor lesions and minimal expression in adjacent normal tissue (**Extended Data Fig. 8c**).

An anti-CA9 nanobody^81^ was recombinantly expressed in *E. coli*, but initial expression yield was low. To optimize expression, we referenced a panel of nanobody stabilizing mutations previously associated with nanobody stability^82^ and identified five residues in the CA9 nanobody sequence commonly found in unstable nanobodies (**Extended Data Fig. 8c**). We used the protein stability prediction tool INPS-MD^83^ to assess the impact of these single-point mutations, each predicted to have stabilizing or neutral effects on free energy (**Extended Data Fig. 8d**). Incorporation of these mutations significantly improved nanobody expression (**Supplementary Fig. S7a, Supplementary Table S8**) while preserving target binding, as confirmed by staining of CT26 lung metastasis tissue with a dye-labeled CA9 nanobody and biodistribution showing increased uptake of the CA9 nanobody in tumor-bearing lungs (**Extended Data Fig. 8e-h, Supplementary Fig. S8**).

To evaluate the biosensors in the CT26 CRC lung metastasis model, mice with and without CT26 lung metastases were administered a cocktail of four biosensors, each incorporating a distinct PSurf-discovered substrate paired with a unique DNA barcode (**Supplementary Fig. S7b**). One hour after administration, urine was collected from each mouse (**Fig. 7c, Extended Data Fig. 9a**). Urinary DNA barcode abundance was quantified using Cas12a-based detection, in which urine samples were incubated with Cas12a and substrate-specific guide RNAs. Cas12a activation, indicating the presence of each DNA barcode, was measured via collateral cleavage of a fluorescent reporter DNA (**Extended Data Fig. 9b)**. Among the biosensors, those incorporating DNLGRAF and PFLYLFG substrates successfully distinguished tumor-bearing mice from healthy controls, as evidenced by elevated Cas12a activation in urine from CT26 tumor-bearing mice (**Fig. 7d**). Receiver operating characteristic (ROC) analysis revealed strong diagnostic performance, with area under the curve (AUC) values ranging from 0.82 to 0.89 for individual sensors, where an AUC of 1.0 represents perfect classification (**Fig. 7e and Extended Data Fig. 9c-e**).

To further enhance detection accuracy, we applied multiple linear regression to integrate signals from all four biosensors into a unified predictive model. Four sensors were selected as previous studies have shown that using 3 to 5 biosensors offers sufficient predictive power in isogenic mouse models, with only slight gains from adding additional sensors^69, 84^. This multiplexed approach further improved classification performance, yielding an AUC of 0.93. These results demonstrate the potential of PSurf-discovered substrate biosensors for noninvasive tumor detection and highlight the value of combining biosensors into a multiplexed panel for robust disease classification.

## Discussion

Dysregulated proteolytic activity in specific disease microenvironments presents compelling opportunities for the development of conditionally activated diagnostic and therapeutic strategies. However, progress in this field has been constrained by the small pool of disease-selective substrate sequences. Most current substrates are discovered through recombinant enzyme assays, often focusing on MMPs and chosen based on transcriptomic data. Yet there is a significant gap between transcriptomic information and actual enzymatic activity in tissues, due to factors such as post-translational modifications, the influence of protease inhibitors, and other regulatory mechanisms affecting these enzymes. Additionally, substrates identified using individual enzymes do not capture the combined activity of the many protease families present in native tissue environments. As a result, directly mapping proteolytic activity and discovering substrates within tissue samples remains an unmet need in protease research.

In this study, we report PSurf, a platform that, to our knowledge, is the first technology to enable direct proteolytic activity mapping and substrate discovery in tissue samples. PSurf builds on traditional yeast surface display methods but incorporates important technological advances, including the design of specific double-tagging strategies and the establishment of differential selection methods (**Supplementary Fig. S9**). These developments were essential for enabling the platform to characterize proteolytic activities in tissue-derived samples.

The success of PSurf with tissue-derived material is attributable to key features of yeast display technology, including compatibility with high-resolution cell sorting, the ability to perform multiple rounds of selection to enrich rare cleavage events even under conditions of low enzyme activity, and the capacity for differential selection strategies to identify context-specific substrates. Moreover, the PSurf library offers much broader substrate coverage than synthetic libraries and is renewable without the high costs of synthesizing new libraries, making it well-suited for broad deployment. Our study thus expands the applications of yeast display technologies beyond their traditional uses^39, 65, 85-90^.

The utility of PSurf was demonstrated through several applications. First, the platform identified novel cathepsin B substrates with varying sensitivities under different pH conditions. These sequences may be applied for designing new antibody-drug conjugate linkers or pH-dependent cathepsin inhibitors. Additionally, PSurf discovered substrates responsive to lung metastases that were more sensitive than widely used MMP-derived probes. These new substrates showed low reactivity in healthy and inflamed lung tissue, indicating their tumor selectivity. Importantly, we developed nanobody-linked fluorogenic biosensors based on these sequences and validated their *in vivo* utility for distinguishing tumor-bearing from healthy animals.

Additionally, this study also provided one of the first global profiles of tissue cleavage, revealing a previously unknown enrichment of phenylalanine and leucine residues in both healthy tissues and tumors. This finding suggests conserved protease cleavage patterns and highlights the need for tissue-based differential selection to identify context-specific probes.

Biochemical analysis of the top tumor-reactive probe further illustrated the power of tissue-based screening. Mass spectrometry identified cleavage sites within the DNLGRAF substrate. Alanine-scanning mutagenesis showed that multiple residues across the seven–amino acid sequence are essential for high proteolytic sensitivity, and that even single amino acid changes can eliminate tumor-specific activity. These results suggest that using a seven–amino acid library, rather than shorter sequences used in many previous methods, can be advantageous for discovering tumor-sensitive probes.

Although the specific proteases responsible for substrate cleavage have not yet been identified, inhibitor studies suggest that multiple enzymes may contribute to the observed activity, highlighting a critical distinction of our selection technology compared to individual enzyme-based screens. Notably, three out of the four nominated probes were not cleaved by recombinant MMPs, indicating that other protease families are likely responsible for their cleavage. These findings demonstrate PSurf’s capacity to identify tissue-responsive substrate sequences without requiring prior knowledge of the specific proteases.

The development of protease-activatable agents, such as antibody-drug conjugates, masked antibodies, masked cytokines, produgs, and activity-based diagnostics, continues to advance across applications in oncology, inflammation, aging, and other medical areas. As these technologies progress toward clinical translation, there is a growing need for strategies like PSurf that enable the discovery of substrates precisely tailored to specific biological contexts. By facilitating the identification of substrates that reflect the true activities within a disease microenvironment, PSurf may provide a powerful platform to support the development of protease-responsive technologies across a broad range of clinical applications.

Overall, PSurf is a broadly applicable platform for tissue-level extracellular proteolysis profiling, substrate discovery, and the development of conditionally activated diagnostic and therapeutic agents. This technology addresses a critical gap in the field of tumor-activated substrate discovery and provides a foundation for novel strategies that harness proteolytic activity as a hallmark of disease. To promote further research and facilitate community-wide application, we will make the PSurf library and associated protocols freely available for non-profit scientific use, contributing an open-access resource to advance the field of protease biology.

